# Simple and Robust Fabrication and Characterization of Conductive Carbonized Nanofibers Loaded with Gold Nanoparticles for Bone Tissue Engineering Applications

**DOI:** 10.1101/2020.03.28.013383

**Authors:** Houra Nekounam, Zahra Allahyari, Shayan Gholizadeh, Esmaeil Mirzaei, Mohammad Ali Shokrgozar, Reza Faridi-Majidi

**Author notes:** Corresponding authors: Reza Faridi-Majidi, Ph.D, Mohammad Ali Shokrgozar, Ph.D.

## Abstract

Bone tissue engineering is a new and applicable emerging approach to repair the bone defects. Electrical conductive scaffolds through a physiologically relevant physical signaling, electrical stimulation, has shown a highly promise in this approach. In this paper, we fabricated carbon nanofiber/gold nanoparticle (CNF/GNP) conductive scaffolds using two distinct methods; blending electrospinning in which GNP were blended with electrospinning solution, and electrospinning/electrospraying in which GNP was electrosprayed simultaneously with electrospinning. The obtained electrospun mats underwent stabilization/carbonization process. The scaffolds were characterized by SEM, XRD, FT-IR and Raman spectroscopy. SEM characterizations showed improved morphology and a slight decrease in the diameter of the spinned and sprayed nanofibers (from 178.66 ± 38.40 nm to 157.94 ± 24.14 nm and 120.81 ± 13.77 nm, respectively), while XRD analysis confirmed the crystal structure of the nanofibers. Raman spectroscopy revealed enhancement in the graphitization of the structure, and the electrical conductivity of the structure improved by up to 29.2% and 81% in electrospraying and blending electrospinning modes, respectively. Indirect MTT and LDH toxicity assays directly were performed to assess MG63 cell toxicity, but no significant toxicity was observed and the scaffolds did not adversely affect cell proliferation. It can be concluded this structure have potential for bone tissue engineering applications.

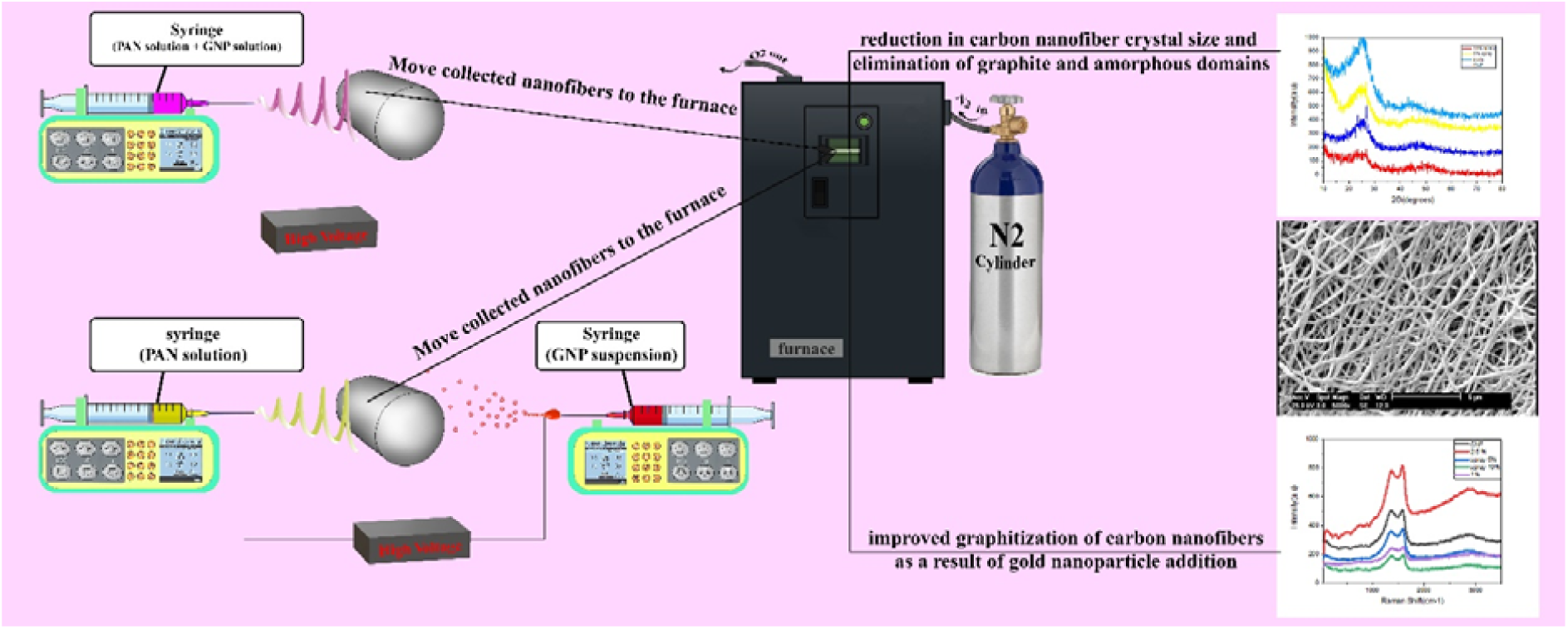

## 1. Introduction

Bone tissue engineering has emerged in recent years as a potential approach to overcome the difficulties in healing, regeneration and functional restoration of bone defects following damages caused by trauma or as a result of osteoporosis amongst other relevant causes.^1–3^ This approach is usually accompanied by the combination of a synthetic, natural or hybrid 2D or 3D scaffolds, cells, and a physiologically relevant chemical or physical signaling.^4–12^

Despite numerous studies focusing on preparation of bone tissue engineering scaffolds and comprehensive investigations of different scaffold component combinations, there is still much-needed improvements necessary for integrating bone tissue engineering scaffolds for in-vivo studies with potential significantly superior clinical outcomes.^13^ Such shortcomings are particularly evident in studies with a major or minor focus on electrical stimulation of bone cells or tissue prior or after implantation.^14,15^

These studies require tailored, engineered and repeatable electrical properties of the scaffolds, while also ensuring high biocompatibility, proper cell attachment, adequate mechanical integrity and tissue integration in addition to the golden missing chain between these properties, which is ensuring reproducibility with low fabrication costs.^16–18^ The latter property is still highly sought despite the development of numerous platforms with significant and promising results.^16,19^

Over the different stages of bone tissue engineering scaffold development, electrospinning have been constantly considered as a suitable fabrication method due to the its perceived multifaceted ability to make nano- and micro-fibrous structures from a variety of polymer solutions with tunable fiber characteristics which have been shown to have a dominant effect on bone cell behavior and tissue integration.^20–24^ Nanofibrous and microfibrous scaffolds have long been used for tissue engineering applications due to their inherent compatibility with native extracellular matrix, their high fabrication yield and overall low fabrication costs.^9^ Electrospinning of conductive polymers have been previously addressed and explored. Common examples of conductive polymers include polypyrrole (PPy), polythiophene and, polyaniline (PANi), with diverse application ranging from biomedical engineering to corrosion protection.^25–28^ Amongst the conductive polymer candidates for electrospun scaffolds, PPy and PANi has been shown to have in vitro and in vivo biocompatibility to various degrees depending on the fabrication parameters and intended cells or tissues, especially yielding successful fabrication and biological results when electrospun in blended format.^29–33^ However, elucidating proper biological response from a composition involving electrospun conductive polymers is challenging despite providing the necessary electrical conductivity, which has led to a noticeable downfall in their application in this field and the biocompatibility of these structures and their precursors is challenging ^34^

Structural and surface modification of electrospun scaffolds have relatively been limited to biological molecule coating, inducing changes in surface roughness and addition of a natural component in the scaffold structure.^35,36^ Despite the improvements obtained through these methods, they add challenges and difficulties to the process, especially in terms of fabrication and reproducibility.^37^

Other potential methods for obtaining electrically conductive scaffolds which have been recently more investigated include addition of conductive nanoparticle components and using standard and reproducible techniques such as furnace-enabled carbonization.^38–40^ Both embedding metallic nanoparticles and carbonization can be considered of high interest for bone tissue engineering, since they are tools for structural and surface modification such as decreasing the contact angle, they also contribute to increasing the electrical activity of the prepared scaffold products and gold nanoparticles (AuNPs) specifically can also be used to induce angiogenesis and differentiation to bone lineage.^41–43^ AuNPs are promising materials for bone tissue engineering due to their easy preparation, tunable chemical and electrical properties, anti-inflammatory properties, and their successful implementation as osteogenic materials for bone tissue regeneration. In addition to their potential as electrically conductive components of scaffolds, it has been shown that they can be used to module osteoblast cell behavior (including differentiation) and to modify pathways responsible for osteoclast formation, indicating that AuNPs are versatile materials for bone tissue engineering applications.^44,45^

Among candidates to be used for bone tissue engineering scaffolds, carbon nanofibers (CNFs) have been shown to be exceptional materials for bone tissue engineering in terms of mechanical properties, tailored surface energy, biologically-relevant nanoscale topography, spatially-controllable bone cell adhesion and bone mineral deposition and biocompatibility. It was demonstrated that CNFs-based bone replacements induce better tissue integration and result in better patient outcomes as compared to traditional commercially available options.^46,47^ Additionally, CNFs possess high electrical conductivity which significantly elevate their potential for conductive scaffold applications.^46^

In the present work, Polyacrylonitrile (PAN) polymer solutions were integrated into conductive nanofibrous scaffolds using electrospinning, followed by carbonization in a tube furnace. Gold nanoparticles were embedded either as a blend in PAN solution or they were co-electrospun with the PAN solution, hence providing flexibility in the fabrication process. *In situ* attachment method was used to fabricate gold nanoparticles on the surface of nanofibers in previous similar works, in which they first synthesized carbon nanofibers and then synthesized gold nanoparticles by adding precursor using different reduction beads or they used the covalent reaction between activated carbon nanofibers and pre-made gold nanoparticles.^48,49^ However, these methods entail certain deficiencies such as complex process, unreliable fabrication and cytotoxicity which create barriers for their in vitro and in vivo applications.^50^ All of the structures that have been developed are mostly based on the role of the sensor and the electrochemical applications with minimal focus on preparing scaffolds with desired biological response and physiologically-relevant electrical cues.

In order to circumvent these issues, a need for simple and cost-effective approaches with minimal potential cytotoxic components is sensed in the literature. Providing electrical conductivity is an integral component of such solution, as it provides a positive physical cue for cell behavior if implemented using safe and simple approaches.^51,52^

As it can be observed in the literature, electrical cues are useful in improving the proliferation and differentiation of various tissues including the nerve, heart, bone and skin. Additionally, precious studies have confirmed the positive effect of gold nanoparticles on bone tissue regeneration. All steps were designed for maximal reproducibility and minimal necessary equipment and efforts. In the first method, gold nanoparticles were added to the prepared polymer solution and then the electrospinning process was performed (blending electrospinning method). In the second method, we used side by side electrospinning systems (with polymer solution on one side electrospun and AuNPs sprayed on the other side).

This was followed by characterizations through scanning electron microscopy (SEM), XRD, FTIR, tensile mechanical testing, electrical conductivity measurements, Raman spectroscopy, contact angle measurements and in vitro biocompatibility assays, and we will compare fabricated scaffolds using the two methods with bare CNFs. We will provide a thorough assessment of applicability of the prepared scaffolds for bone tissue engineering and electrical stimulation applications, and we will show, for the first time, that this structure is shown to be nontoxic and may be appropriate for bone tissue engineering applications.

## 2. Experimental

### 2.1 Materials

Polyacrylonitrile (PAN, MW=80000), Dimethylformamide (DMF), sodium citrate, and HAuCl_4_ were purchased from Merck (Germany). Fetal Bovine Serum (FBS), RPMI 1640 without phenol red and DMEM-high glucose media were all purchased from Gibco (USA). MTT Formazan was purchased from Sigma-Aldrich. Cytotoxicity Detection KitPLUS (LDH) was purchased from Roche. Human osteosarcoma cells, MG-63, were supplied by the National Cell Bank of Iran (NCBI), Pasteur Institute of Iran (NCBI, C555).

### 2.2 Nanofibrous scaffold synthesis and carbonization

#### 2.2.1 Preparation of carbon nanofibers

To provide spinning polymer solution, we dissolved PAN powder in DMF solution at 60 °C under magnetic stirrer for 12 hours to prepare 9 wt% polymer solution of PAN. The electrospinning processes were carried out using an electrospinning equipment (Electroris, FNM, Tehran, Iran).

The electrospinning was carried out by applying a voltage of 20 kV between the needle and the collector with rotation speed of 400 rpm and rate of 1 mm/hr at room temperature. Nanofiber plates electrospined with a thickness 100-120 µm.

The electrospun mats was peeled from the aluminum foil and was treated with heat to stabilize and carbonize nanofibers to obtain carbon nanofibers. Stabilization and carbonization of PAN electrospun mats was done in a tube furnace (Azar, TF5/25-1720, Iran) according to our previous study.^46^ To prevent the PAN electrospun mats from shrinkage during the stabilization, PAN nanofibrous mats were cut into rectangular shape and pasted on graphite block by PAN solution. Stabilization was performed in air at 290 °C with heating rate of 1.5°C/min and holding at 290 °C for 3h. The stabilized nanofibers were then taken apart from the graphite blocks and were carbonized at 1000°C under high purity nitrogen atmosphere (N2 99.9999%, Air Products). The samples were heated at rate of 4 °C/min and kept for 1 hr at 1000 °C.

#### 2.2.2 Gold nanoparticles/CNF blend electrospinning method (GCNF Blend)

Three concentrations of gold nanoparticles were blended in 7% PAN solution. The polymer concentration was reduced from 9% w/v to 7% w/v due to droplet formation during electrospinning for higher concentrations, which prevented effective electrospinning. 7% PAN polymer solution was first prepared, since gold nanoparticles did not allow PAN polymerization DMF solution. The solution was then stirred at 60 ° C and 50 ppm gold nanoparticle in water solution was later added with 1 w/v%, 2.5 w/v% and 5 w/v%. After stirring for 5 hr, the solution was electrospinned with 22 V, at a 10 cm distance, with a rate of 1 mm/hr to obtain gold nanoparticle-containing PAN nanofibers. Prior to this, gold nanoparticles were synthesized using a modified Turkevich method for gold nanoparticle synthesis to obtain 49 ppm gold nanoparticles with 25 nm size, which were then centrifuged at 10000 rpm for 20 min to obtain higher concentrations.^53^

#### 2.2.3 Gold Nanoparticles/CNF electrosprayed method (GCNF Sprayed)

In the electrospraying method, polymer solution and gold nanoparticles were electrospun separately. Alternatively, 5% and 10% gold nanoparticles solutions (which were then centrifuged at 10000 rpm for 20 min to obtain higher concentrations) were co-electrospinned with PAN nanofibers. This fabrication method relies on a side by side electrospinning systems, with one nozzle for nanofiber electrospinning, while the other nozzle is used for gold nanoparticle electrospraying. Stabilization is done similar to previous method, but the carbonization temperature is reduced by 800 °C under previous conditions.

### 2.3 Scanning Electron Microscopy

Scanning Electron Microscopy (SEM) (CM200-FEG-Philips) was used to characterize the micro- and nanostructure of the scaffolds. The mages were obtained using an acceleration voltage 25 kV and a working distance of 13.6 mm. Energy dispersive X-ray (EDX) was used to characterize dispersion of gold nanoparticles in the scaffold structure.

### 2.4 Electrical conductivity measurement

A simple four point apparatus was used for measuring the resistivity of semiconductor samples. Similar to previous work,^54^ passing a current through two outer probes and measuring the voltage through the inner probes allows the measurement of the substrate resistivity. Five samples were tested for each condition.

### 2.5 Raman spectroscopy

Raman spectroscopy was conducted to assess the crystallinity of the carbonized structures. This method has proved to be effective for characterizing carbon-based structures. D-band and G-band of the spectra represent graphitic carbon and structural disorder near the edge of crystalline structure, respectively, and the ratio of peak intensity is generally used for characterizing structural disorder. G-band appears near 1582 cm^-1^, while D-band is found around 1350 cm^-1^.^55–57^ Raman spectrum of the samples were performed using Teksan Takram P50C0R10 raman spectrometer (laser wavelength: 532 nm, laser power: 0.5-70 mW).

### 2.6 Fourier-transform infrared spectroscopy (FTIR)

The IR spectrum obtained from FTIR spectrometer lies in the mid-IR region between 4000 and 666 cm^-1^. Transition energies corresponding to changes in vibrational energy state for many functional groups are located in the mid-IR region and they can be indicative of appearance of an absorption band in this region can be used to determine whether specific functional groups exist within the structure. This characterization method was used to assess the structural changes in the samples due to the addition of gold nanoparticles.^58^

### 2.7 X-ray diffraction

A high resolution X-ray Diffractometer (PANalytical X’Pert Pro) with a Cu Kα source (λ = 0.1540598 nm) was used to analyze the crystallinity of carbonized nanofibers and to investigate their microscopic structure.^59^

### 2.8 Contact angle measurement

Contact angle measurement was performed to assess the hydrophilicity of the surface. As described elsewhere, a contact angle of less than 90° denotes a hydrophilic surface, while a surface with a contact angle of more than 90° can be considered as a hydrophobic surface.^60^

### 2.9 MTT assay

Due to absorbance of MTT dye on carbonized nanofibers, indirect MTT assays were performed for the scaffolds. As described elsewhere, 1 ml cell culture media without FBS per 0.1 gr scaffold was added to each sample and the incubated cell culture media was collected after 24, 48 and 72 hr and supplemented with 10% FBS, and were subsequently added to wells containing MG-63 cells. Alternatively, for the direct cell culture on the scaffolds, they were sterilized with 70% ethanol and UV, and were placed and fixed on the bottom of a 48-well plate. Fixation using medical grade O-rings (C. Otto Gehrckens GmbH & Co, Germany) was necessary to avoid any movement and to ensure proper cell seeding on top of the scaffolds. Cell seeding densities of 7000, 5000 and 3000 cell/cm2 was used for 24, 48 and 72 hr experiments, respectively.

### 2.10 LDH Cytotoxicity assay

Lactate dehydrogenase (LDH) assay was first used to assess the cytotoxicity of our scaffolds. LDH Cytotoxicity kit (Roche Diagnostics, Germany) was used for this test according to the company protocol using negative and positive controls according to following formula:

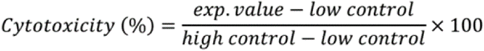

Similar to previous section, the scaffolds were rinsed with %70 ethanol and UV sterilized. MG-63 cells were cultured on the scaffolds seeding densities of 7000, 5000 and 3000 cell/cm2 was used for 24, 48 and 72 hr experiments, respectively. RPMI media without phenol red and 1% FBS was used due to interference of phenol red and FBS with LDH adsorption.

### 2.11 LDH proliferation assay

Identical conditions were used for cell seeding on the scaffolds were placed in a 37 °C incubator with 90% humidity and 5% CO2 for 24, 48 and 72 hr. Cell culture media was extracted after the incubation periods and 100 μl fresh DMEM media was added to each well. Lysing solution was added to each well and incubated for 15 min to ensure complete cell lysing. The entire cell culture media was transferred to other wells and mixed with LDH reaction mixture. Absorbance was measured at 290 nm after 10 min in room temperature. Proliferation was calculated using the following formula:

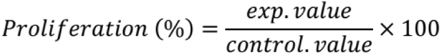

### 2.12 Cell attachment and morphology

Scanning electron microscopy (SEM) was used to assess cell attachment and morphology on CNF and CNF/2.5% AuNP scaffolds. Briefly, MG-63 cells were seeded on both types of scaffolds and they were fixed with 2 v/v% glutaraldehyde. This was followed by consecutive steps of 20%, 40%, 60%, 80%, and 96% ethanol treatment, each for 10 min. The samples were sputter-coated with gold and SEM was used for visualization of cell attachment and morphology.

### 2.13 Statistical analysis

The experiments were performed with at least 5 repeats and the results were expressed as means with standard deviation. One-way analysis of variance (ANOVA) was used and P<0.05 was used as the level of significance for statistical analysis.

## 3. Results and discussion

### 3.1 Energy dispersive X-ray and Scanning Electron Microscopy

EDX and SEM results of the scaffolds are shown in figure 1 and 2. EDX was performed to map the gold nanoparticles inside the scaffold structure and as it can be observed, contrary to what could have been a plausible issue, even after exposure to 800°C did not deteriorate the stability of gold nanoparticles inside the nanofibers and an almost homogenous distribution of gold nanoparticles is observed throughout the structure. Although cell morphology in SEM images is improved in the presence of 1% gold nanoparticles, the nanofibers are thick and are merges together due to the furnace heat. However, higher concentrations of gold nanoparticles contributed to separation of the fibers. As opposed to 5% concentration which exhibited more interfiber attachment and thicker fibers, sprayed samples and 2.5% concentration led to a reduction in nanofiber diameter, which can be attributed to gold nanoparticle surface charge.

**Figure 1.**
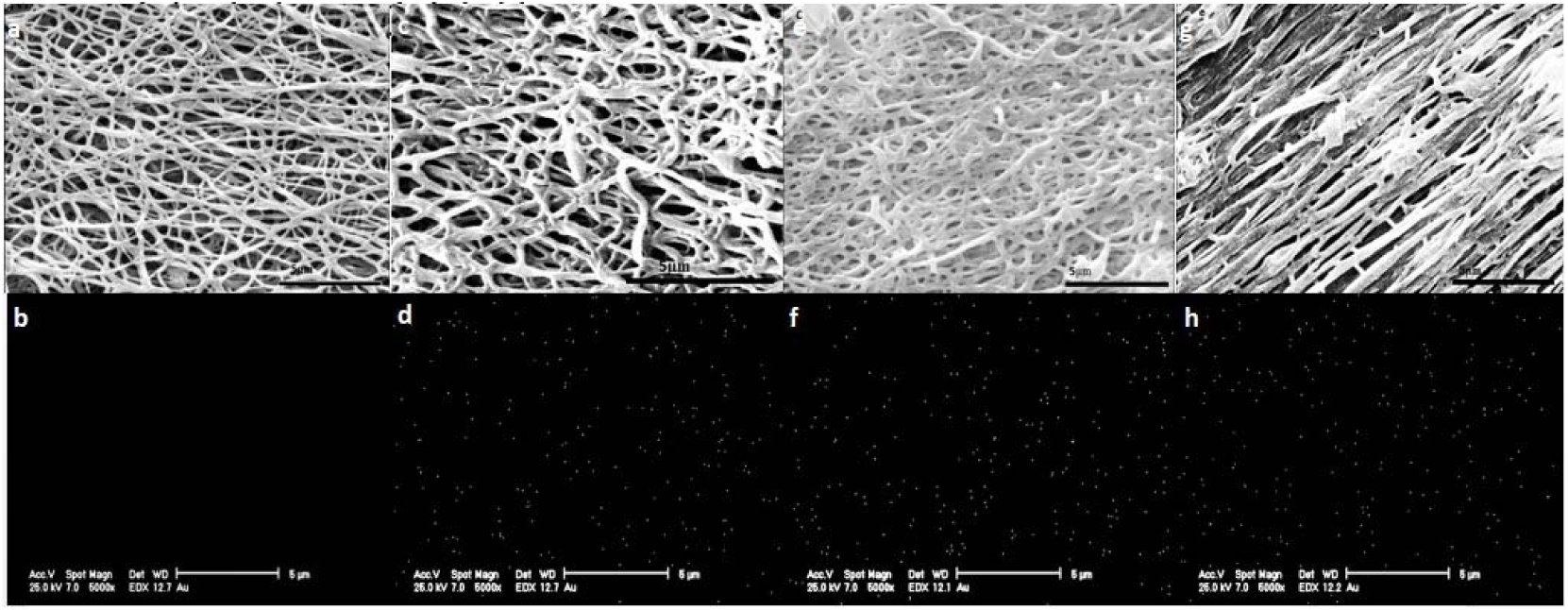
SEM and EDX characterization of the electrospinnned scaffolds. a) SEM of CNF, b) EDX of CNF (Au) c) SEM of CNF/1% AuNP, d) EDX of CNF/1% AuNP, e) SEM of CNF/2.5% AuNP f) EDX of CNF/2.5% AuNP(Au) g:SEM of CNF/5%AuNP, h) EDX of CNF/5% AuNP(Au)

**Figure 2.**
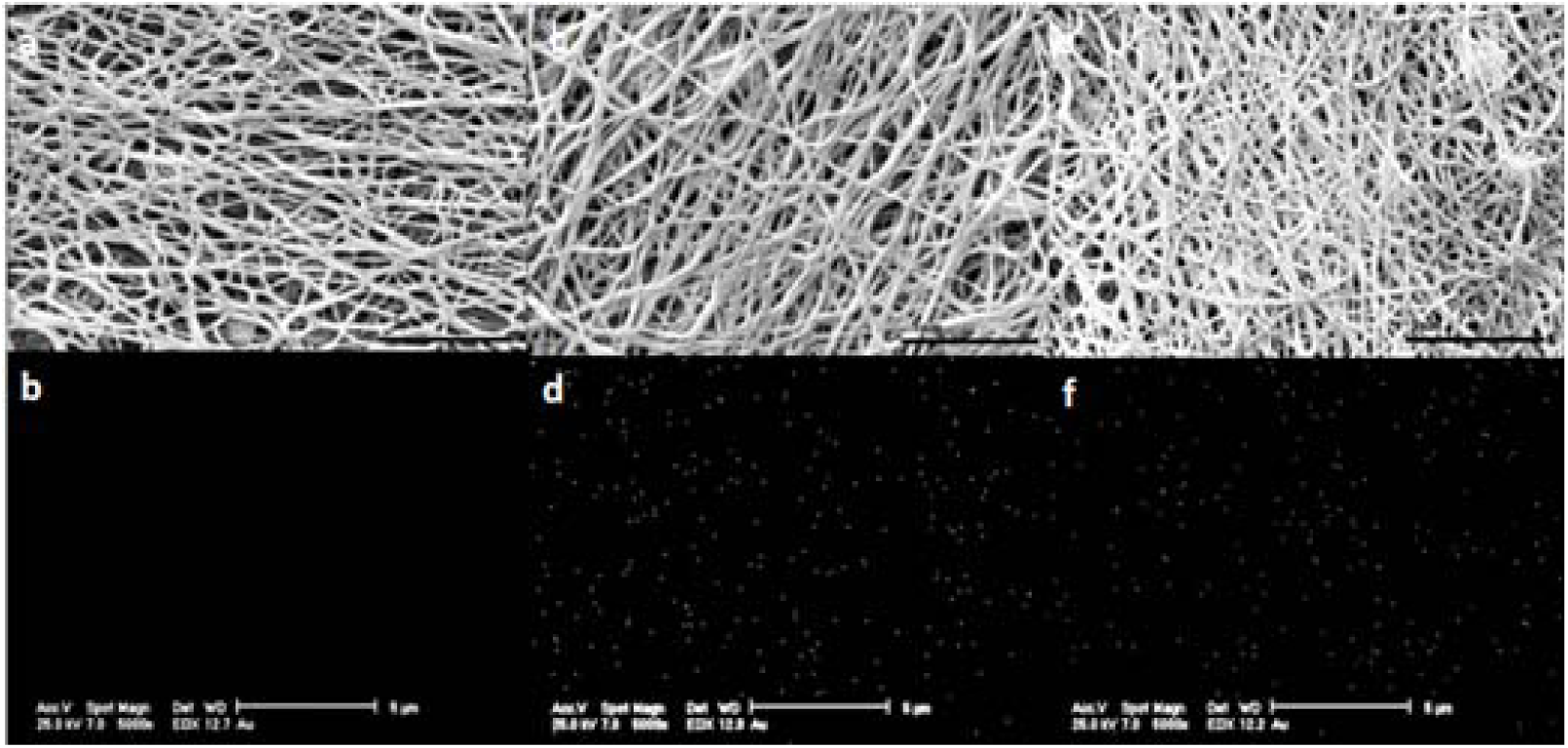
SEM and EDX characterization of the electrosprayed scaffolds. a) SEM of CNF, b) EDX of CNF(Au), SEM of CNF/5% AuNP sprayed, d) EDX of CNF/5% AuNP sprayed, e) SEM of CNF/10% AuNP sprayed, f) EDX of CNF/10% AuNP sprayed(Au).

As observed in figure 2, EDX results demonstrate a significant decrease of gold nanoparticle presence in electrosprayed form. This occurred despite the higher concentration of introduced gold nanoparticles. This can be attributed to the entrapment of gold nanoparticles in between the fibers in the mixture format as opposed to the loss in sprayed format. Electrosprayed scaffolds barely show any presence of gold nanoparticles and are almost indistinguishable from PAN fibers with no gold nanoparticles. Size-distribution of nanofibrous scaffolds has been presented in Table 1, which shows consistent fiber size distribution across all sample despite the addition of gold nanoparticles.

**Table 1.** Nanofibrous scaffold fiber sizes under different conditions.

| Sample |  | Fiber size (nm) | Sample |  | Fiber size (nm) | Sample |  | Fiber size (nm) |
| --- | --- | --- | --- | --- | --- | --- | --- | --- |
| PAN | | $152.90 \pm 24.51$ | 2.5% AuNP Spinned | Before furnace | $122.54 \pm 29.23$ | 5% AuNP Sprayed | Before furnace | $178.83 \pm 29.10$ |
| CNF | | $178.66 \pm 38.40$ | | After furnace | $157.94 \pm 24.14$ | | After furnace | $120.81 \pm 13.77$ |
| 1% AuNP Spinned | Before furnace | $165.73 \pm 26.87$ | 5% AuNP Spinned | Before furnace | $140.15 \pm 29.69$ | 10% AuNP Sprayed | Before furnace | $177.16 \pm 27.93$ |
| | After furnace | $182.56 \pm 52.22$ | | After furnace | $259.27 \pm 51.73$ | | After furnace | $157.8 \pm 24.58$ |

These results collectively show the effectiveness of our method for homogenous distribution of nanoparticles. As opposed to previous works which experience much difficulty in homogenous dispersion of gold nanoparticles, our simple and robust method prevented nanoparticle agglomeration.^61–63^ An alternative method in the letter based on adding gold nanoparticles after carbon nanofiber synthesis is also associated problems such as difficult integration of nanoparticles in the structure, which was avoided with our technique.

### 3.2 Electrical conductivity measurement

The results of electrical conductivity measurements have been presented in Table 2. There is a statistically significant increase in conductivity upon addition of 2.5% gold nanoparticles. However, addition of 5% gold nanoparticles led to a significant decrease in electrical conductivity, suggesting a possible disintegration in the nanofiber structure, leading to an adverse effect on conductivity.

**Table 2.** Electrical properties of Carbonized Nanofibers under different conditions.

| Charachterisics of the samples | 1%AuNP | 2.5%AuNP | 5%AuNP | 5% Sprayed | 10% Sprayed | CNF |
| --- | --- | --- | --- | --- | --- | --- |
| Conductivity(s.cm<sup>-1</sup>) | 3.15±0.32 | 4.96±0.06 | 2.65±0.06 | 3.36±0.25 | 3.54±0.18 | 2.74±0.02 |
| Resistivity(Ω.cm) | 0.33±0.093 | 0.25±0.056 | 0.37±0.01 | 0.29±0.031 | 0.28±0.082 | 0.36±0.027 |

### 3.3 X-ray diffraction

X-ray diffraction (XRD) analysis revealed amplified peaks at 2Θ=26° and 2Θ=44° corresponding to (001) and (002) planes, respectively (Figure 3). Additionally, the present weak peaks corresponding to gold nanoparticles confirmed their presence in the fibers for the mixture mode and on the surface for the sprayed mode. Peaks corresponding to gold nanoparticles were observed with different intensities in 2Θ=38°, 2Θ=44°, 2Θ=64°, and 2Θ=77° for (111), (200), (220), and (311) planes, respectively (Table 3).

**Table 3.** XRD data of CNF and CNF/AuNP.

| Samples | 2θ | B (radian) | D(002) ($\text{\AA}^\circ$) | Lc (nm) |
| --- | --- | --- | --- | --- |
| CNF | 26.52 | 0.19 | 3.36 | 70 |
| CNF/1% AuNP(Blend) | 28.45 | 0.49 | 3.13 | 20 |
| CNF/2.5% AuNP(Blend) | 26.22 | 0.78 | 3.66 | 14 |
| CNF/5% AuNP (sprayed) | 26.65 | 0.29 | 3.34 | 46.66 |
| CNF/10% AuNP (sprayed) | 26.62 | 0.19 | 3.35 | 70 |

**Figure 3.**
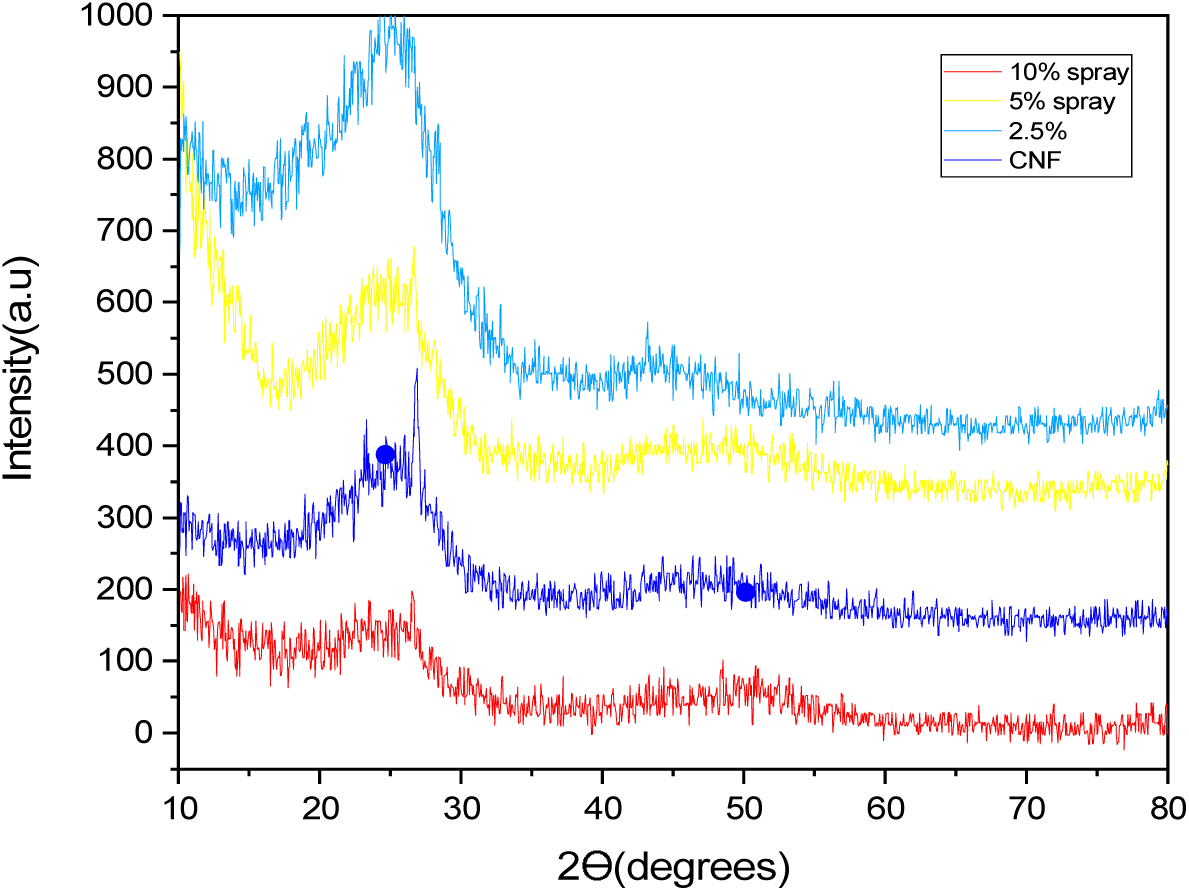
The X-ray diffraction (XRD) pattern of CNF and different combinations of CNF/AuNP.

These results verified that using 800 °C furnace temperature didn’t deteriorate the structural integrity of gold nanoparticles, which is consistent with previous reports of around 1000 °C melting temperature for nanoscale gold nanoparticles.^61^

XRD results confirmed the presence of graphite nanocrystals and shows a reduction in carbon nanofiber crystal size and elimination of graphite and amorphous domains. Smaller domains with graphite structure are also present in the samples. Wider peaks (especially in 2.5% samples) imply more crystalized structure. XRD analysis for scaffolds with mixture format showed the best performance in terms of graphite crystal.

Additionally, XRD software analysis confirmed the presence of gold nanoparticles in 10% spray mode more distinctively more than the mixed mode samples. This can be attributed to the necessity of certain percentage to be detected by the tool (5% for mixed mode). One additional possibility for mixed mode samples is the presence of more gold nanoparticles in the inner layers of nanofibers. However, since the gold nanoparticles are on the surface and are in high concentration in the spray mode, they were easily detected by the XRD tool. The results showed the presence of gold nanoparticle crystals in cubic form with a=4.06 and b=4.06 and the carbon structures in hexagonal form with a=2.45, b=2.45 and c=6.69, collectively confirming the high thermal properties of gold nanoparticles in our scaffolds.

### 3.4 Raman spectroscopy

Raman spectroscopy characterizations exhibited a decreased ID/IG ratio in samples with gold nanoparticles (Figure 4 and table 4). This was particularly evident in 2.5 % gold nanoparticle samples, which confirmed the improved graphitization of carbon nanofibers as a result of gold nanoparticle addition. The gradual decrease in G values is a result of a decrease in graphite crystal size and elimination of amorphous carbon, which is consistent with XRD results. This can be attributed to surface plasmon resonance phenomenon of gold nanoparticles creating localized heat which has potentially contributed to an improvement in crystallinity and the graphite structure. D/G ratio corresponds to edge to basal plane in graphite crystals and can be representative of graphite domains. ID¬/IG ratios confirm an increase in graphite to edge ratio and a decrease in amorphous domains.

**Table 4.** Peak Position and relative ratio of Raman peak intensities of CNF and CNF/AuNP.

| Samples | D position (cm<sup>-1</sup>) | G position (cm<sup>-1</sup>) | R(I<sub>D</sub>/I<sub>G</sub>) |
| --- | --- | --- | --- |
| CNF | 1372 | 1582 | 97% |
| CNF/1% AuNP(Blend) | 1376 | 1602 | 97% |
| CNF/2.5% AuNP(Blend) | 1376 | 1566 | 89% |
| CNF/5% AuNP(Blend) | 1355 | 1576 | 100% |
| CNF/5% AuNP (sprayed) | 1361 | 1552 | 92% |
| CNF/10% AuNP (sprayed) | 1372 | 1597 | 91% |

**Figure 4.**
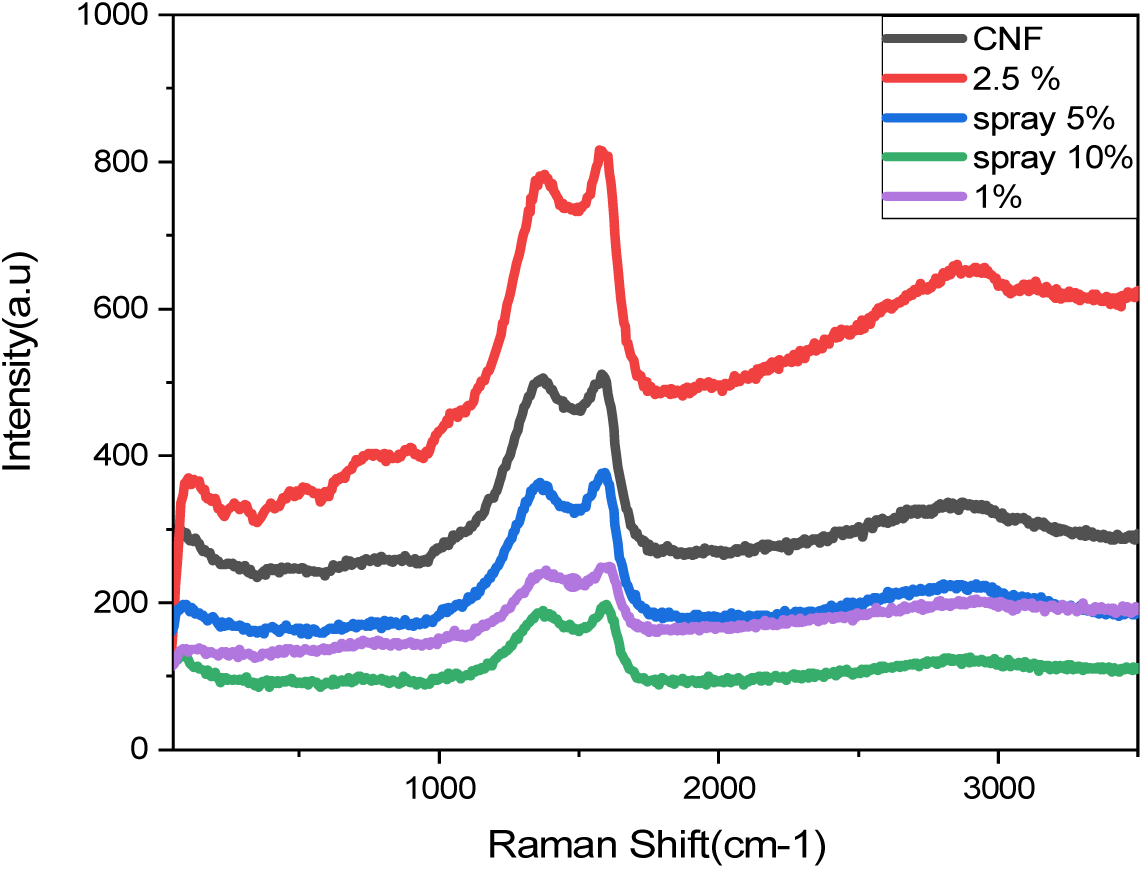
Raman Spectra of CNF and CNF/AuNP.

### 3.4 Fourier-transform infrared spectroscopy

Fourier-transform infrared spectroscopy (FTIR) showed sharp peaks between 1800 and 1900 cm-1 wavelengths (Figure 5). These peaks are not present in fibers containing 2.5% gold nanoparticles, which confirms atomic level changes in carbon nanofiber surface due to the presence of gold nanoparticles.

**Figure 5.**
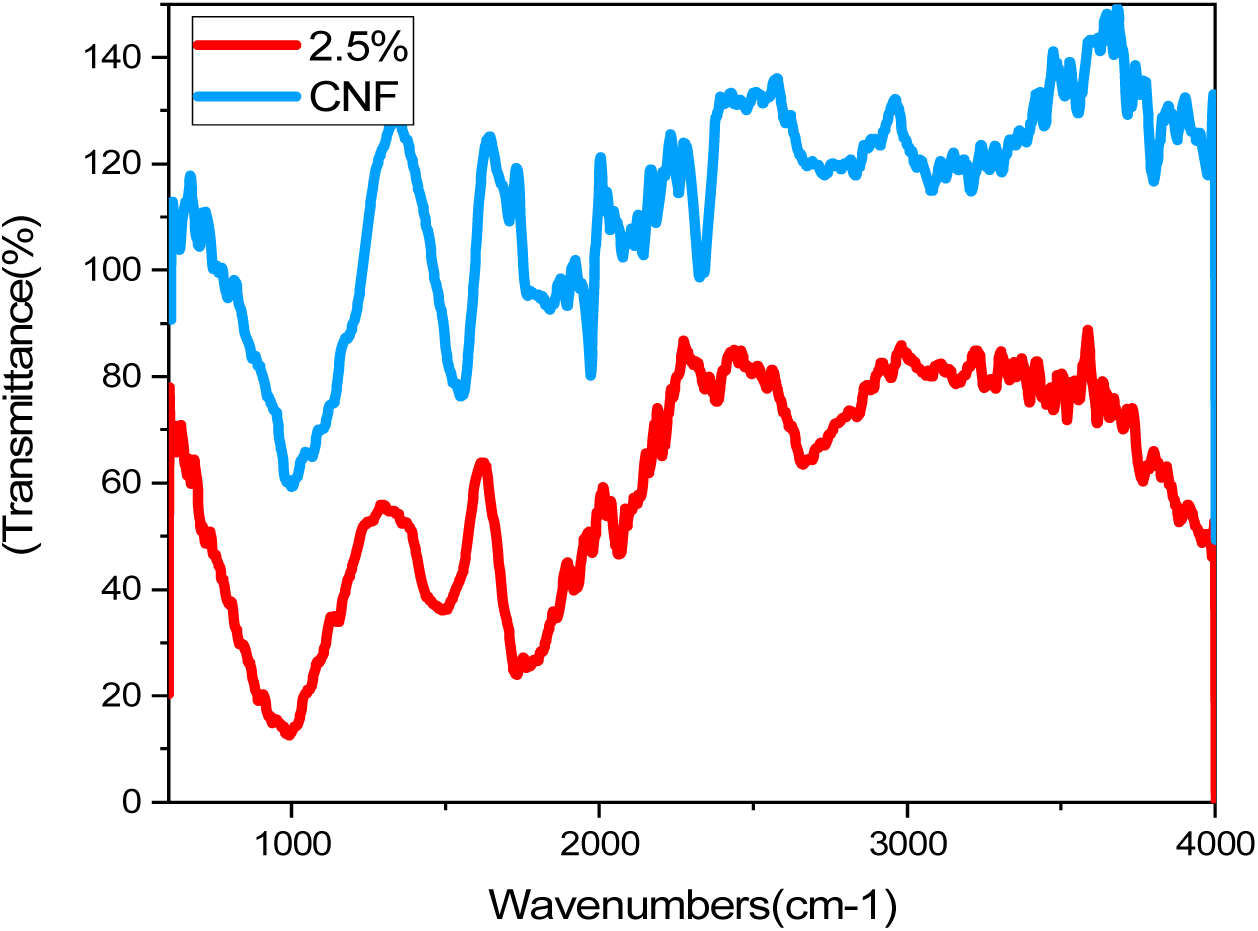
FT-IR Spectra of CNF and CNF/AuNP.

### 3.5 Contact angle measurement

Contact angle measuements did not reveal any changes on the contact angle upon addition of gold nanoparticles and the surface hydrophilicity remained unchanged (Figure 6).

**Figure 6.**
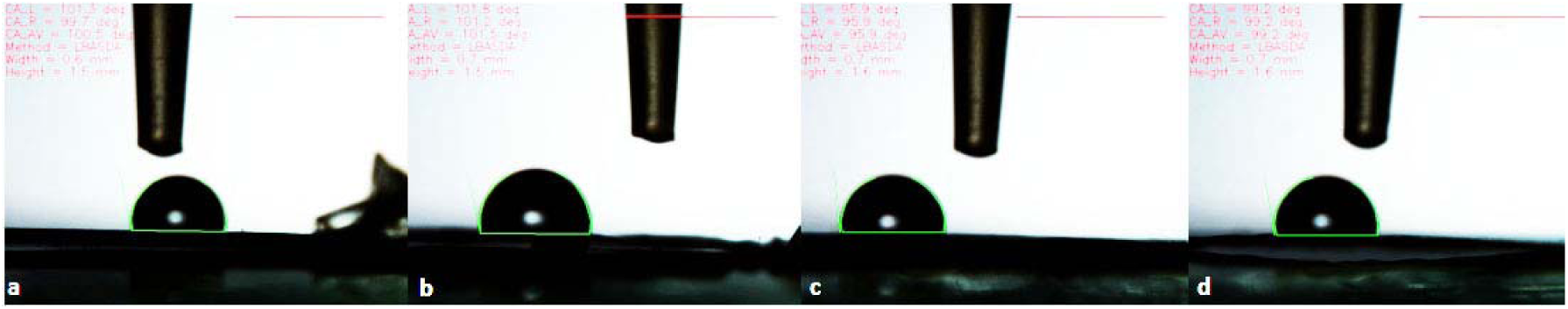
Contact angle measurement of nanofibrous scaffolds. a) CNF, b) PAN/CNF, c) CNF/ 2.5% AuNP, CNF/ 5% sprayed AuNP.

### 3.6 MTT assay

As stated before, indirect MTT assay was performed to assess the potential cytotoxicity of the prepared scaffolds. Although there was a slight decrease in cell viability in the prepared scaffolds and especially in the electrosprayed samples, this decrease was not statistically significant and therefore, the scaffolds all possessed biocompatibility (Figure 7). This argument is further solidified, considering that prolonged exposure (72 hours) shrunk this difference. Previous attempts on integration of gold nanoparticles in carbon-based structures were mostly focused towards application of such structures as electrodes, but here we demonstrate the efficacy of our design for biological application as evidenced by MTT, cytotoxicity and proliferation assays.^64–66^ These results are superior or at least similar as compared to previous works that have attempted to fabricate bone tissue scaffolds with similar compositions.

**Figure 7.**
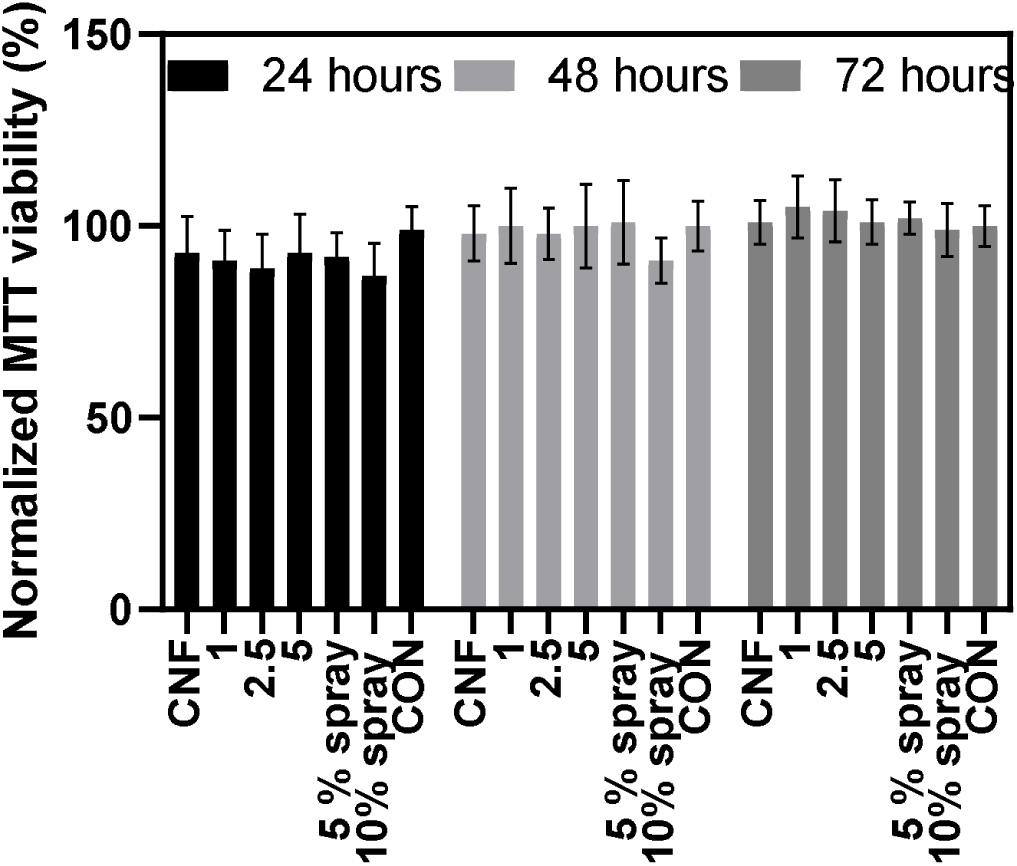
The indirect MTT assay result on the cell viability of Mg63 cells.

### 3.7 LDH Cytotoxicity assay

LDH cytotoxicity was utilized as an orthogonal method to assess scaffold biocompatibility. Although there was some perceived cytotoxicity over different time periods, still even in the worst case, cytotoxicity did not exceed %10, which is acceptable considering the components of the scaffolds and a simple comparison with previous reports further confirms potential superiority of our scaffold fabrication and modification as opposed to previously established methods (Figure 8).^61,67,68^

**Figure 8.**
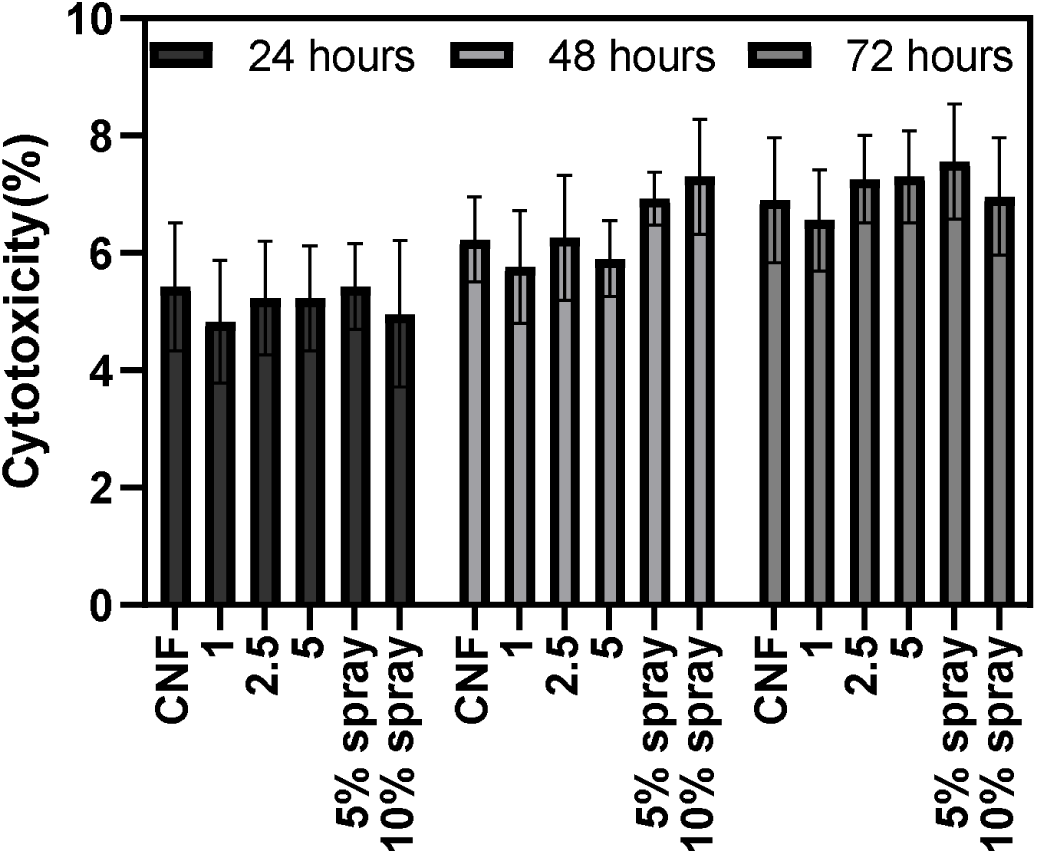
The LDH results of cytotoxicity on Mg63 Cells.

### 3.8 LDH proliferation assay

In addition to no significant cytotoxicity, any suitable scaffold should facilitate cell growth and proliferation as compared to golden standards such as tissue culture plastic controls (Figure 9). LDH proliferation assay showed significant proliferation on the scaffolds, comparable to tissue culture plastic controls. This suggests that cell seeding and subsequent proliferation on the prepared scaffolds can potentially be used to create cell-laden scaffolds for in vitro or in vivo assays which could pave the way for clinical applications of the prepared scaffolds.

**Figure 9.**
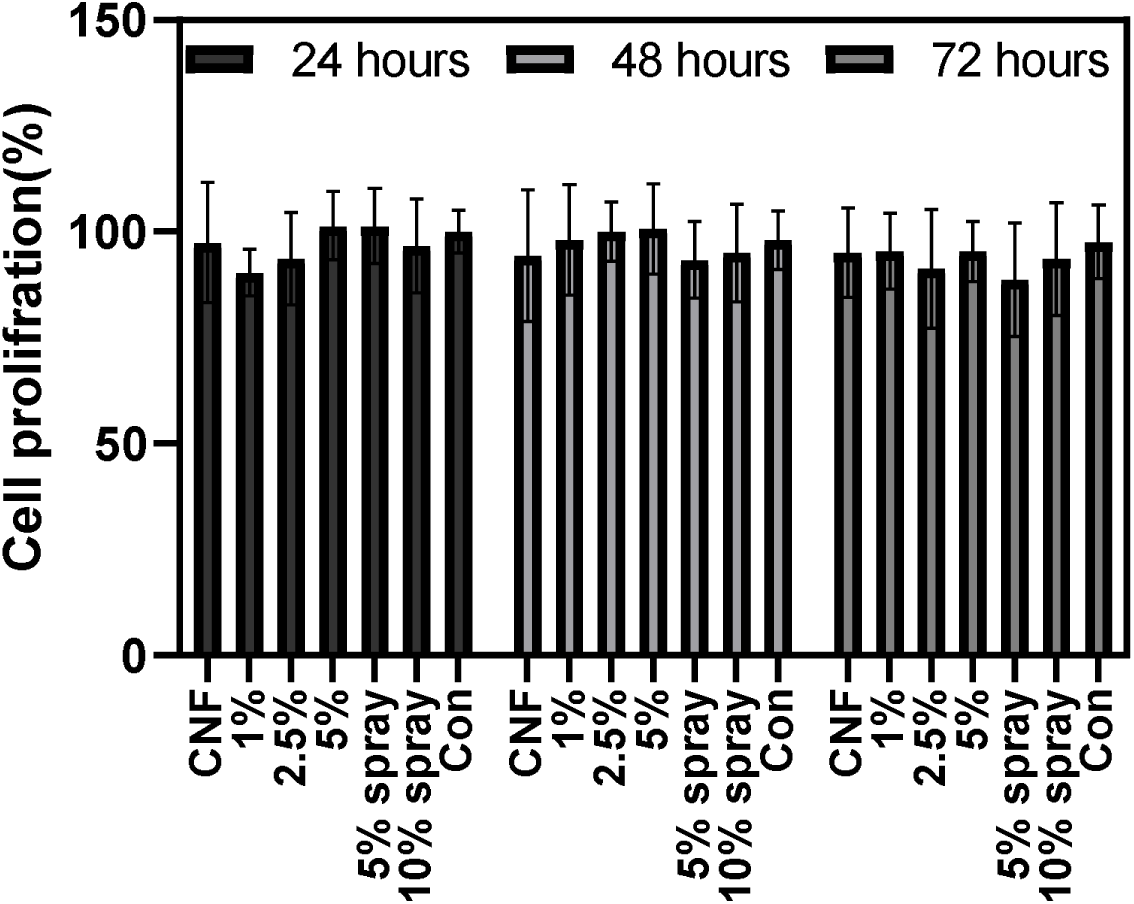
Cell Proliferation of Mg-63 Cells on CNF and CNF/AuNP.

### 3.9 Cell attachment and morphology

As discussed previously, cell culture, fixation and scanning electron microscopy (SEM) was performed for a preliminary assessment of cell attachment and morphology. As it can be observed from Figure 10, SEM showed proper cell attachment and cell spreading both on carbon nanofibrous scaffolds with and without the presence of gold nanoparticles. Normal morphology and attachment of MG-63 cells can be interpreted as promising with future continual of this work for further *in vitro* and *in vivo* studies as the use of similar structures in the literature has entailed controversy regarding their potential cytotoxicity. Considering these results and previous data shown in this work on direct and indirect cell cytotoxicity, we have obtained a method with minimal cytotoxicity as opposed to similar works in the literature.

**Figure 10.**
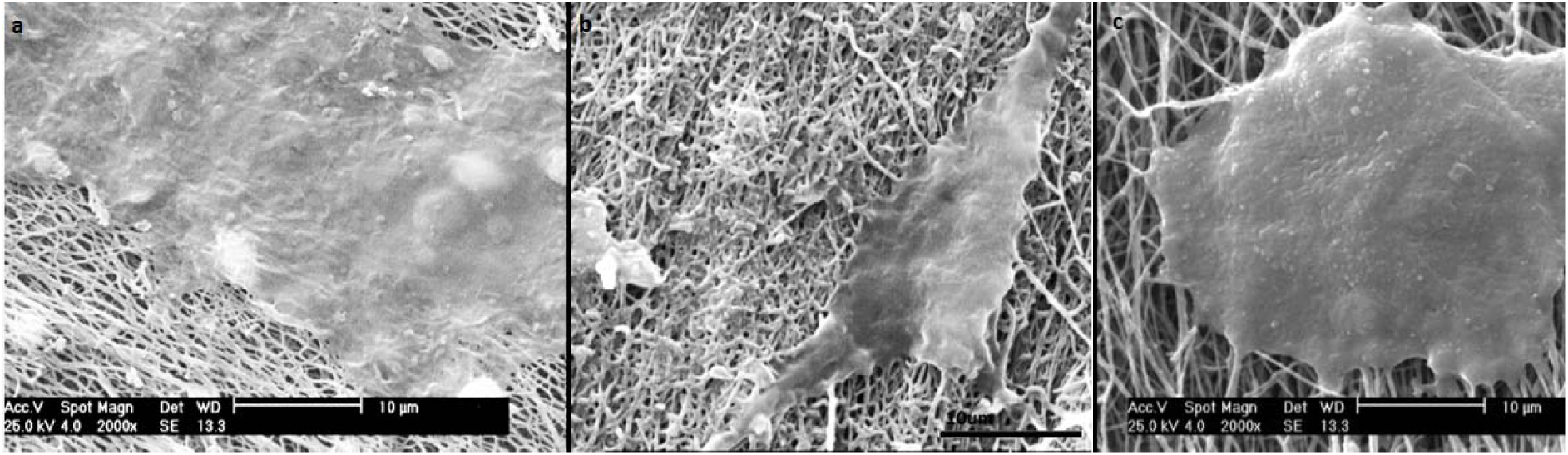
Cell attachment and morphology on CNF (a) and CNF/2.5% AUNP scaffolds (b) 5% GNP Sprayed on CNF(c)

## 4. Conclusions

In this work, through simple and highly controllable fabrication methods and by relying on incorporation of gold nanoparticles in a carbon nanofiber matrix, not only we improved the electrical and physical properties of the scaffolds, we were also able to maintain and significantly improve the biological response and cell-substrate interactions of our scaffolds.

We observed that gold nanoparticles can be embedded in carbon nanofibers structures using two distinct methods (GCNF blended and GCNF sprayed). The effects of gold nanoparticles on the carbon nanofibers structure were investigated through comprehensive characterizations. The addition of nanoparticles in both methods resulted in a decrease nanofibers sizes but was more effective in the electro-spraying format, in which uniform size distribution and a better morphology with less adhesion between nanofibers was observed. This may be attributed to electrostatic repulsion induced by gold nanoparticles on the surface of nanofibers. The XRD results demonstrated that improvement was observed in carbon nanofibers crystals in the presence of gold nanoparticles. On the other hand, the crystalline form of the gold nanoparticles was shown to indicate that the furnace process had no effect on the structure of the gold nanoparticles. Raman spectroscopy results showed graphitic structure enhancement that probably was caused due to local heat induced by the plasmon surface of the nanoparticles and this could explain the improvement in the electrical conductivity of the structure. The FT-IR results also showed changes in the surface chemistry of the nanofibers, but the changes were not sufficient to result in changes in the surface hydrophilicity, considering contact angle results show no significant difference between samples containing gold nanoparticles and carbon nanofibers. Toxicity tests indirect MTT and LDH showed that the carbon nanofibers and nanofibers containing gold nanoparticles (GCNF blended and GCNF sprayed) did not induce any cytotoxicity and was biocompatible with the MG-63 cells, while the results of cell proliferation showed no difference between the control and carbon nanofibers and GCNF blended and sprayed.

Such scaffold fabrication strategy can be further explored in future studies in combination with exogenous in vitro electrical stimulation, with potential expansion to in vivo studies and eventually reaching clinical work. However, such translation requires further biological and biophysical characterization and more in-depth research into the underlying mechanisms for any improvement in regeneration as a result of electrical conductivity and electrical stimulation. We believe that the presented results provide suitable platforms for such endeavors in our research group and also, in the tissue engineering community.

Most of previous applications for scaffolds with similar formulations in the literature were for detection and sensor applications and did not thoroughly assess cytotoxicity or proliferation, in contrast with our current study which thoroughly addressed cytotoxicity and proliferation for bone cells.^64–66^ Our results have potentials and implications for future work on osteogenic differentiation and assessing the in vitro and in vivo effects of electrical conductivity and exogenous electrical signaling, stemming partly from successful embedment of gold nanoparticles.

